# Active tactile discrimination is coupled with and modulated by the cardiac cycle

**DOI:** 10.1101/2022.02.10.479959

**Authors:** Alejandro Galvez-Pol, Pavandeep Virdee, Javier Villacampa, James M Kilner

## Abstract

Perception and cognition are modulated by the phase of the cardiac signal in which the stimuli are presented. This has been shown by locking the presentation of stimuli to distinct cardiac phases. However, in everyday life sensory information is not presented in this phase-locked and passive manner, instead we actively sample the world. Whether active sensing is coupled and modulated with the cardiac cycle remains largely unknown. Here we recorded the ECGs of human participants while they actively performed a tactile grating orientation task. Here we show that the duration of subjects’ touch varied as a function of the cardiac phase in which they initiated it. Touches initiated in the systolic phase of the cardiac cycle were held for longer periods of time than touches initiated in the diastolic phase. This effect was driven by the elongation of their holds to sample the most difficult gratings. Conversely, while touches in the control condition were coupled to the cardiac cycle, their length was not modulated as a function of when in the cycle these were initiated. In line with interoceptive inference accounts, these results are consistent with the hypotheses that we actively adjust our sensory sampling so that we spend more time in the diastole period of the cardiac cycle in which perceptual sensory sensitivity is greatest.

## Introduction

Over half a century ago, it was proposed that the sense of touch is built upon the processing of our own body in space and the stimuli in our environment (exteroception and proprioception; Gibson, 1962). Yet, today we know that the nervous system also senses, interprets, and integrates signals originating from within the body (i.e., interoception; Craig, 2002; Khalsa et al., 2018). Importantly, research on interoception has shown that the signals from within the body do not carry information about the state of the body, but also influence how we process exteroceptive sensory information and stimuli (Al et al., 2020; Azevedo et al., 2017; Azzalini et al., 2019; Critchley & Garfinkel, 2018; Garfinkel et al., 2014; Leganes-Fonteneau et al., 2020; Salomon et al., 2016).

The most studied source of interoceptive signals is the heart. As an intrinsic oscillator, the heart has two phases: in the *systole* phase the heart contracts and ejects the blood, whereas in the *diastole* phase it expands while being filled. Both phases comprise a full cycle. Previous studies have shown that participants’ responses to stimuli vary according to the phase of the heartbeat in which exteroceptive stimuli are presented. Perceptual sensitivity to painful, visual, tactile, and auditory stimuli are typically reduced when they are presented in systole compared with diastole (Al et al., 2020; Edwards et al., 2001, 2007; Grund et al., 2022; McIntyre et al., 2006; Motyka et al., 2019; Pramme et al., 2016). It has been proposed that these cardiac-related sensory attenuation effects reflect the competing allocation of attentional resources (Berntson & Khalsa, 2021; Critchley & Garfinkel, 2018; Khalsa et al., 2018) during periods in which there are afferent signals from baroreceptors (Critchley & Harrison, 2013; Garfinkel et al., 2014). Within this framework, it is proposed that there are periods in the cardiac signal in which participants are more sensitive to exteroceptive signals (diastole) and periods in which participants’ processing is less sensitive (systole) due to concurrent cardiac related afferent signals.

The effects of the cardiac cycle upon exteroceptive processing has been demonstrated by the presentation of brief stimuli to participants timed to occur during their systole or diastole phases. However, in our everyday lives, exteroceptive stimuli are unlikely to be presented to us passively in a cardiac phase dependent manner. Here and in our previous work, we have adopted a more ecological approach in which participants actively move to seek information as they wish (i.e., active sensing). Indeed, we have recently shown that in an active sampling visual paradigm, saccades and visual fixations are coupled to the systole and diastole phases of the cardiac cycle (Galvez-Pol et al., 2020). These results are consistent with the hypothesis that interoceptive and exteroceptive processing adjust to each other by sampling the environment during the most quiescent periods of the cardiac cycle (visual fixations during diastole). In the present study, we expand upon this and other work in active vision (Kunzendorf et al., 2019; Ohl et al., 2016) to test the prediction that when we move to touch objects we do so in a manner that reflects the relative perceptual sensitives of touching in systole and diastole.

While active touch sensing has been studied in animals and humans (Grant et al., 2014; Olczak et al., 2018; Prescott et al., 2011; Schaefer et al., 2009; Vega-Bermudez et al., 1991), no accounts of interoceptive signalling originating from the heart have been considered. Here we tested the hypothesis that the timing of active touch sensing would be coupled to the distinct phases of the cardiac cycle, and that such coupling would be linked to changes in participants’ responses. To test this, we instructed our participants to perform an active tactile discrimination task of grating orientation. We co-registered the initiation, stationary holds (to sense the gratings), the end of participants’ touches, and their electrocardiograms (ECGs). For each touch, we computed the phase of the cardiac cycle in which each tactile event occurred, as well as the concomitant consequences of such an alignment in participants’ responses. From an interoceptive inference perspective, studies in passive sensing have shown that tactile sensitivity is reduced during the systole phase (Al, Iliopoulos, et al., 2021; Grund et al., 2022; Motyka et al., 2019), therefore, we predicted that participants behaviour would reflect this and would sample more in the diastole phase than systole.

## Materials and methods

### Participants

Psychophysiological studies and analyses of cardio-sensory processing usually involve sample sizes on the order of 30-45 participants (Al et al., 2020; Galvez-Pol et al., 2020; Grund et al., 2022; Herman & Tsakiris, 2021; Kunzendorf et al., 2019; Motyka et al., 2019). Accordingly, we recruited 50 healthy adults (32 females; age range = 18-35) who participated in the current study. All participants reported normal cardiac condition, volunteered to take part in the experiment, gave informed consent, and were reimbursed for the participation. Ethical approval for the study was obtained from the UCL research ethics committee. Two participants were excluded due to a faulty recording of their responses, and two more due to performance close to chance level across all levels of difficulty. In the control condition, one more participant was excluded from the analysis due to a faulty recording of the ECG data. Thus, the data of 46 participants were included in the analysis of the main experimental condition, and 45 in the control condition; these conditions were analysed separately.

### Task and procedure

Participants were seated with the forearm of their dominant hand rested on the top of a table in palm up position. Participants took part in an active touch sensing paradigm where they were presented with tactile gratings of different widths (narrow to wider gratings; see Stimuli and apparatus). From the participants’ perspective, half of these gratings were horizontal and the other half vertical. The gratings were randomly presented through a custom-built device by the experimenter, who was seated opposite to the participants. The participants’ task was to touch the gratings (one per trial) with the index finger of the dominant hand to determine their orientation. To do this, they had to move their index finger up, make contact with the grating, move their finger down and respond verbally to state the orientation. Then, the experimenter keyed the response. Importantly, once the grating was positioned, the participants were free to initiate, hold, and finalise the touch (i.e., they started, held, and ended touching when they felt like). Therefore, sensation arose through their movement rather than through the passive movement of the gratings.

Before the experiment, the participants received instructions (see Instructions in Supp. materials) and completed one practice block of 14 trials. If they did not have further questions, they completed 12 more blocks of 14 trials while their ECG and responses were recorded. In the last two blocks, no grating was presented and instead a flat stimulus of the same diameter was used. The participants were instructed to follow the same instructions, but with no need to report any orientation at the end of the trials. This was the movement control stimulus.

### Stimuli and apparatus

We used a custom-built device to present the gratings to the participants. This had two parts: a small platform with a rail, and a sliding section that held fifteen gratings (14 gratings with seven widths by two orientations, and one flat stimulus). The later section could slide along the former and allow presenting one grating at the time through an opening in the centre of the device, above the participants’ index finger. The sliding section was moved by the experimenter, who presented the gratings in random order by following a list generated in Matlab R2016b (The MathWorks Inc., Natick, MA, USA). The device, gratings, and list of gratings to-be-presented were only visible to the experimenter. After each trial, the experimenter keyed the participants’ verbal responses in a computer.

The stimuli were made using the Ultimaker-2 3D printer (Ultimaker, Geldermalsen, NL) and were designed using online modelling software (Tinkercad; Autodesk Inc, California, US). The gratings consisted of two copies of seven gratings with increasing width of 0.4mm (from 0.412mm to 2.8mm). Each copy was positioned in the device either vertically or horizontally. When the participants made contact with these stimuli, a continuous pulse at 1000Hz was sent to a CED Power Unit 1401-3A. This signal was recorded simultaneously with the participants’ ECG in Spike2 8.10 (Cambridge Electronic Design Limited, Cambridge, UK). The amplitude of the continuous pulse was set according to the width of the presented grating. Therefore, the pulse itself contained the data required to compute the onset, duration, difficulty, and offset of participants’ touches.

### Data preprocessing and ECG recording

The ECG was recorded using a D360 8-Channel amplifier (Digitimer Ltd., Hertfordshire, UK) in Spike2 8.10; sampling rate 1000 Hz. The ECG electrodes (Skintact Fannin Ltd, Dublin, IE) were placed over the right clavicle and left iliac crest according to Einthoven’s triangular arrangement. To investigate whether participants’ touch varied along the cardiac cycle, as well as along the cardiac phases comprised in the cycle (systole and diastole), we proceed to identify the start and end of each phase. The start of each cycle and the systole phase was detected by computing the R-peaks of the QRS complex in the ECG. To this aim, we filtered the ECG (3-30Hz) using the HEPLAB toolbox implemented in EEGLAB (Delorme & Makeig, 2004; Perakakis, 2019, respectively). We computed the local maxima of the ECG by using the *findpeaks* function in Matlab with a minimum peak to peak distance of 550 ms. By doing so, we obtained the timepoints of the R-peaks occurring during participants’ touches. Next, we proceed to identify the end of the systole phase (start of the diastole phase) for each cardiac cycle. The change of phase can be estimated by computing the end of the T-wave in the ECG. This was done by locating the T-peak as a local maximum after the QRS complex and calculating the area of a series of trapeziums along the descending part of the T-wave. The point at which the trapeziums’ area is maximal approaches the end of the systole phase in the ECG (Vázquez-Seisdedos et al., 2011). We visually inspected the R-peaks and end of the systoles returned by the algorithm and applied small adjustments when required. While it is true that the length of the electrical systole in the ECG cannot be fully equated with the mechanical systole (Fridericia, 1921), both are closely tied under normal conditions (Fridericia, 1921; Boudoulas et al., 1981; Gill and Hoffmann, 2010; Motyka et al., 2019).

Trials in which the participants made consecutive contacts (i.e., more than one touch per grating and trial), responded verbally to state the orientation of the grating before finalising the touch, held their touch less than 100ms, more than 5000ms, or were ±3SD of the individuals’ mean duration, were excluded from further analysis. Likewise, given the temporal consistency between R-peaks and end of T-waves, trials in which this interval was separated by ±1.5 × median of the participants’ interval in milliseconds were not considered; in total 94.5% of the total number of trials were kept for further analyses.

## Data analysis

### Touch responses as a function of absolute cardiac cycle

To examine whether there was a significant statistical relationship between when participants touched and the cardiac cycle, we analysed the data in two different ways. First, circular statistics were employed to exploit the repeating nature of the cardiac cycle. We calculated the phase of each event of interest (touch initiation, mean contact phase of the touch, and end of touch) as a function of the whole cardiac cycle. For instance, for an R-R interval of time t_R,_ where the start of a touch occurred at time t_E_, we calculated t_E_/T_R_ × 360. This results in values between 0° and 360° for each event (0 indicating the start of the cardiac cycle). Then, we calculated the participants’ mean phase for each event. As in previous research, we tested separately whether the participants’ mean phase differed from a uniform distribution using Rayleigh tests (Al et al., 2020; Alejandro Galvez-Pol, McConnell, et al., 2020; Kunzendorf et al., 2019; Motyka et al., 2019; Ohl et al., 2016). We performed this analysis for the start, mean of the touch, and end of all touches, separately for the gratings and the control flat stimulus. This analysis was subsequently performed for each level of difficulty.

### Touch responses as a function of cardiac phases

The previous analysis takes into account the repeating nature of the cardiac cycle, but it does not consider its biphasic nature. Previous studies have shown that responses to stimuli vary as a function of the phase of the cardiac cycle in which information is processed (i.e., systole and diastole; e.g., Garfinkel et al., 2014; Al et al., 2020; Leganes-Fonteneau et al., 2020; Grund et al., 2021). In a second analysis, we examined participants’ responses as a function of the phase of the cardiac cycle, systole or diastole. To this end, we defined systole as the time between the R-peak and the end of the T-wave. We used the systolic length of each cardiac cycle to define diastole as a window of equal length, which we located at the end of the cardiac cycle (Motyka et al., 2019; Al et al., 2020). The identical length of these windows was used to equate the probability of having an event in the two phases of the cardiac cycle. To determine the end of t-wave, a trapezoidal area algorithm was applied in each trial (see Data preprocessing). This method has advantages compared to an approach with fixed windows (e.g., defining systole as the 300-ms time window following the R-peak) because it accounts for within- and between-subject differences in the length of systole and diastole.

We analysed i) the proportion of events (start and end touches) occurring during the systolic and diastolic windows, and ii) the duration of touches as a function of the cardiac phase in which they were initiated. We computed these analyses through rmANOVA and follow-up T-tests for all touches, as well as separately for each level of difficulty according to the gratings’ width. Mauchly’s W was computed to check for violations of sphericity, Greenhouse– Geisser adjustments to the degrees of freedom were applied when appropriate, and P values were corrected for multiple comparisons using the Holm-Bonferroni method.

### Data availability and transparency statement

We report *t* and *p*-values, mean and standard deviations, 95% confidence intervals, and effect sizes. We used in-house and standard code to pre-process and plot our data. Specifically, the pre-processing code was implemented in Matlab R2018a (The MathWorks Inc., Natick, MA, USA), the values for analyses and plots were computed in JASP (JASP Team 2021, Version 0.16) and in R using the *circular* package (Version 0.4-93). The code, anonymised data, as well as the resulting analyses are available in the corresponding Open Science Framework repository: https://osf.io/d7x3g/. We report how we determined our sample size, all data exclusions, manipulations, instructions given to the participants, and measures in the study (see Materials and Methods, and Supp. materials). No part of the study procedures was pre-registered before the research was conducted.

## Results

### Descriptive statistics

The proportion of rejected trials (see Methods) comprised 5.5% of the total number of trials, SD = 4%. The mean number of computed heartbeats during participants’ touch was 540, SD = ±173. Participants’ mean heart rate was 76 bpm, with a mean time between R-peaks of 793 ms and SD = ± 110 ms. The percentage of correct responses was 82.7%, SD ± 6.6%, and the mean duration of participants’ touch in the whole experiment was 1103ms, SD = 457.40 ms, Mdn = 999 ms. The mean duration of the systole phase was 342 ms and the SD = 23.4 ms, which is the expected duration for the mean heart rate of our sample (Mao et al., 2003).

### Inferential statistics

#### Proportion of correct responses and holding time as a function of gratings difficulty

In an initial analysis we tested whether the participants’ performance was modulated by the task difficulty. To this end, we compared the proportion of correct responses for each level of difficulty (7 levels). The results of the rmANOVA showed a significant effect of task difficulty on participants’ performance (*F*(3.340, 150.279) = 154.3, *p* < 0.0001, *η*_*p*_^*2*^ = 0.774). Post hoc comparisons showed that participants’ performance for the first four levels of difficulty was fairly similar (all *p* > 0.086; See Supp. Table 1 for all Post hoc comparisons). Yet, performance in these levels was significantly greater than in the remaining levels of difficulty (all *p* < 0.0001, *d* > 0.833). Likewise, the proportion of correct responses in difficulty level 5 was significantly greater than those in difficulty levels six and seven of (both *p* < 0.0001, *d* > 1.618; Figure 2A).

**Figure 1.**
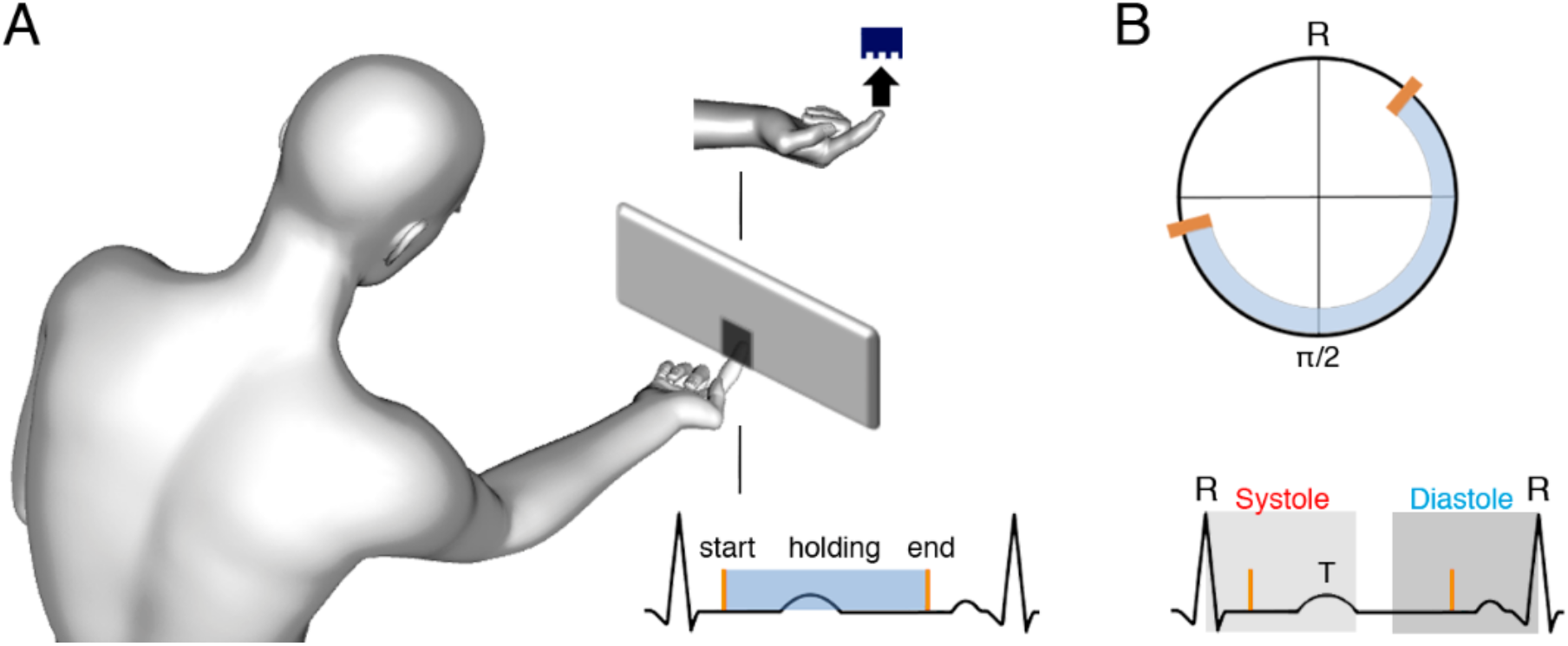
Task and schematic illustration of one trial with cardiac and tactile events. **(A)** Participants’ task was to touch gratings (one per trial) with the index finger to determine their orientation (vertical or horizontal, p = 0.5). Participants were free to start, hold, and end the touch when they felt like while their ECGs were coregistered. **(B)** Upper panel: for each touch, we computed the start, stationary hold, and end of touch in degrees relative to the entailing heartbeat: e.g., for an R-R interval of time tR, where the touch started at time tE, we calculated tE/TR x 360. Then, each type of event was subjected to circular averaging. Lower panel: we also computed the proportion of touches starting and ending in the systole and diastole phases of the cardiac cycle. We used the systolic length of each cardiac cycle to define a diastolic window of equal length (to equate events’ probability), which we located at the end of the cardiac cycle.

**Figure 2.**
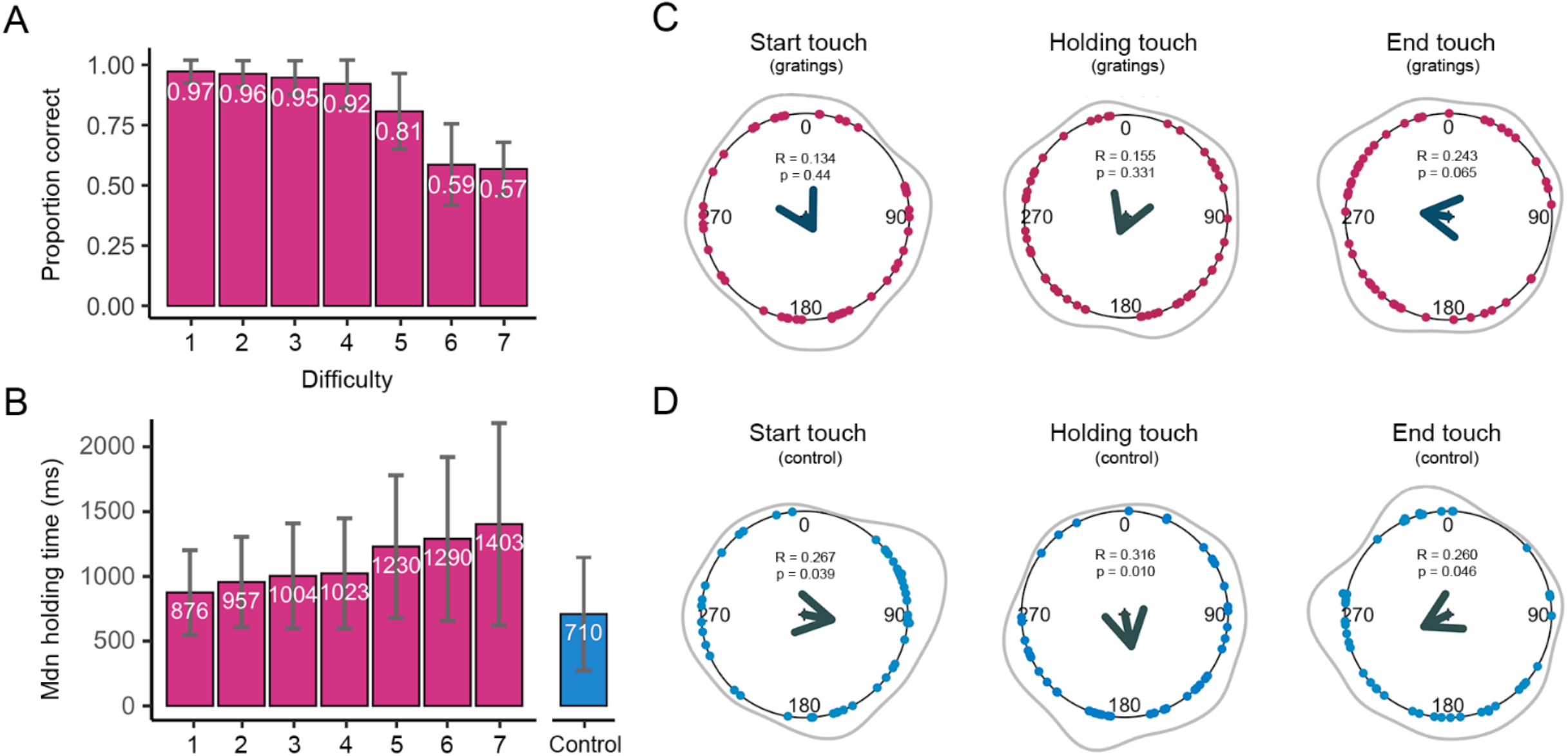
Tactile responses and touch events as a function of the entire cardiac cycle. **(A)** The mean proportion of correct responses changed as a function of task difficulty. Performance in the first four levels of difficulty was similar and significantly greater than in the remaining levels of difficulty; lines show SDs. **(B)** The duration of participants’ stationary holds for the first four levels of difficulty was fairly similar and significantly shorter than in the remaining levels of difficulty. Participants’ stationary holds for the control flat stimulus was significantly shorter (*vs*. all levels difficulty). **(C)** Distribution of starting, holding, and ending touches for the grating stimuli across the cardiac cycle (interval between two R-peaks at 0/360°). Points depict subjects’ mean degrees, the central arrows point to the overall mean degree and its length indicates the coherence of individual means. The grey outer lines depict the circular density of individual means. Overall, the start, mean point of stationary hold, and end of touches occurred at 151°, 202°, and 278° (*p values* = 0.44, 0.331, 0.065, respectively). **(D)** Touch data for the control flat stimulus as analysed and depicted in panel C. Here, the start, mean point of holding, and end of touches occurred at 101°, 168°, and 243° (*p values* = 0.039, 0.010, 0.046, respectively).

Next, we examined the length of time that participants spent holding their touch for each level of difficulty, i.e., time sensing gratings with different widths. Here we took the median of participants’ holding touch in milliseconds. There was a significant effect of task difficulty on the time spent holding the touch (*F*(1.703, 76.641) = 25.13, *p* < 0.0001, *η*_*p*_^*2*^ = 0.358). Post hoc comparisons showed that the length of holding time for the first four levels of difficulty was similar (all *p* > 0.065), as well as shorter than in the remaining levels of difficulty (all *p* > 0.01, *d* > -0.416; see Supp. Table 2 for all Post hoc comparisons). Lastly, we compared the length of holding time touching all gratings and the flat stimulus, where no discrimination was required. The results show that the holding time for the control flat stimulus was significantly shorter than for the gratings across all seven levels of difficulty (*p* ≤ 0.05, *d* > - 0.416; see median times of holding touch Figure 2B).

#### Start, holding, and end of touches as a function of the entire cardiac cycle

Here we analysed the coupling of the participants’ responses (i.e., start, mean contact point of the holding touch, and end of touch) with their own heartbeats. We computed the circular phase of these tactile events as a function of the whole cardiac cycle. Then, we used circular averaging for all trials, as well as separately for each level of difficulty.

In an initial analysis, we pooled data from touches across all difficulty levels. The start of touches occurred on average during the first half of the cardiac cycle at 151°. The phase of the initiation of touches in circular space did not differ significantly from a uniform distribution (Rayleigh test z = 0.134, *p* = 0.44; Figure 2C). The mean phase of the holding touch occurred on average between heartbeats at 202° and its distribution did not differ from uniformity (Rayleigh test z = 0.155, *p* = 0.331). Likewise, the distribution of ending touches in circular space, which occurred on average during the latter part of the cycle at 278°, did not differ significantly from a uniform distribution (Rayleigh test z = 0.243, *p* = 0.065). Subsequently, we analyzed all trials separately for each level of difficulty. While some of the analyses seemed to yield a significant result, they did not reach significance after correcting for multiple comparisons (using the Holm-Bonferroni method for 7 estimates).

Finally, we examined the same events (start, mean contact point of holding, and end of touches) when the participants were instructed to touch the control flat stimulus. Thus, no orientation judgment was required, only movement. The start of touches for the flat stimulus occurred on average during the first half of the heartbeats at 101° and the distribution of these touches differed significantly from a uniform distribution (Rayleigh test z = 0.267, *p* = 0.039; Figure 2D). Regarding the holding of touches, these occurred on average during the mid-part of the heartbeats at 168° of the cardiac cycle. The distribution of these touches in circular space differed significantly from a uniform distribution (Rayleigh test z = 0.316, *p* = 0.010). Last, the end of touches occurred on average during the second half of the heartbeats at 243°. The distribution of these touches was marginally different from a uniform distribution (Rayleigh test z = 0.260, *p* = 0.046).

Overall, these results indicate that when only movement was required, participants’ touches were coupled with the cardiac cycle. These results are consistent with previous findings in active vision (e.g., Kunzendorf et al., 2019; Galvez-Pol et al., 2020) where the movement to sample and sampling of stimuli that did not require discrimination were coupled to the early and mid-phases of the cardiac cycle. Yet, while the present analysis considers the repeating nature of the cardiac cycle, it does not reflect its biphasic physiological nature (contraction and expansion, i.e., systole and diastole phases) nor differences in how participants performed the task.

#### Start and end of touches as a function of cardiac phases

Here we analysed the proportion of starting and ending touches in the systole and diastole phases. For the grating stimuli, we performed a rmANOVA to compare the proportion of events in these cardiac phases across the seven levels of difficulty. For the start of touches, there were no significant main effects of cardiac phase (*F*(1, 45) = 0.194, *p* = 0.662, *ηp2* = 0.004, and task difficulty *F*(6, 270) = 5.535e^-31^, *p* = 1.000, *ηp2* = 3.285e^-14^). Likewise, the interaction between cardiac phase and difficulty was not significant (*F*(6, 270) = 1.123, *p* = 0.349, *ηp2* = 0.024). In contrast, for the end of touches there was a significant main effect of cardiac phase (*F*(1, 45) = 6.334, *p* = 0.015, *ηp2* = 0.123). The proportion of touches ending in systole 0.48 (SD = 0.049) was significantly smaller than in diastole 0.52 (SD = 0.049), (*t*(45) = -2.760, *p* = 0.008, *d* = - 0.407, 95% CI mean difference [-0.065, -0.007]). The effect of difficulty (*F*(6, 270) = 5.670e^-31^, *p* = 1.000, *ηp2* = 2.848e^-14^), and the interaction between cardiac phase and task difficulty were not significant (*F*(6, 270) = 0.688, *p* = 0.659, *ηp2* = 0.015; see Figure 3A,B).

**Figure 3.**
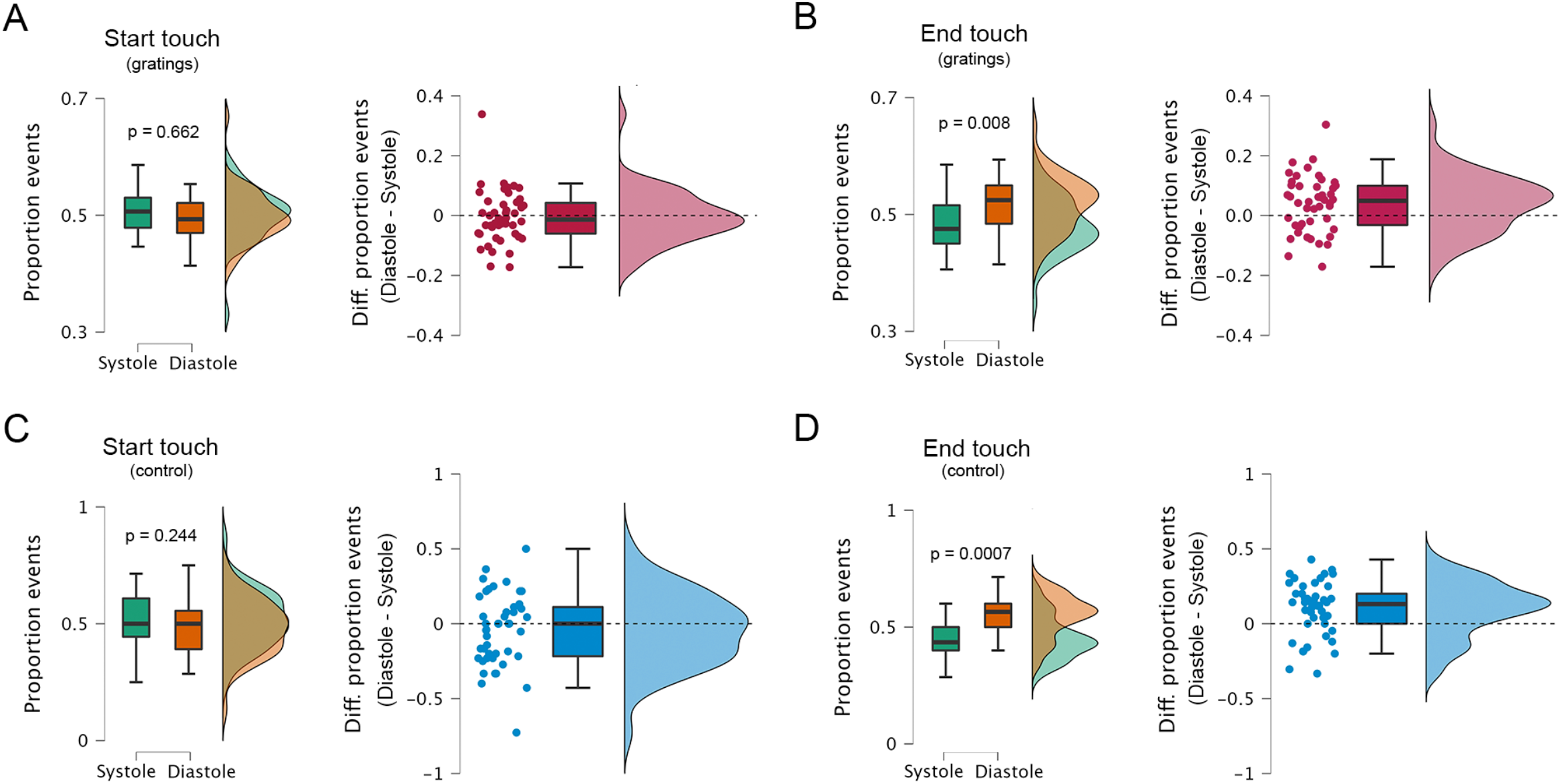
Proportion of starting and ending of touches as a function of cardiac phases. **(A)** For the grating stimuli, the proportion of starting touches was similar in the systole and diastole phases of the cardiac cycle (*p* = 0.710). **(B)** Conversely, the proportion of ending touches was significantly different in the systole and diastole phases, with more touches ending in the diastole phase (*p* = 0.008). **(C)** For the control flat stimulus, the proportion of starting touches was also similar in the systole and diastole phases (*p* = 0.254). **(D)** For the ending, the proportion was significantly different in the systole and diastole phases, with more touches ending in diastole (*p* = 0.0006). Right panels show the difference between events occurring in diastole and systole (i.e., for each participant, proportion of diastole events minus systole events).

For the control flat stimulus, we performed paired sample T-test to compare the proportion of touches starting or ending in the systole and diastole phases. For the start of touches, the results indicate a not significant difference between cardiac phases (*t*(44) = 1.181, *p* = 0.244, *d* = 0.176). Conversely, for the end of touches there was a significant difference between cardiac phases. The proportion of touches ending in systole 0.45 (SD = 0.09) was significantly smaller than in diastole 0.55 (SD = 0.09), *(t*(44) = -3.671, *p* = 0.0007, *d* = -0.547, 95% CI mean difference [-0.157, -0.046]. See Figure 3C,D). These results mirror and expand those found in the above-mentioned circular analysis. Here, participants did not initiate the sampling of information in a particular phase of the cardiac cycle, but were more likely to finish in diastole, which has been reported as the quiescent period of baroreceptor activity and other concurrent physiological fluctuations (Constantinos, 2002; Critchley & Harrison, 2013).

#### Time sensing as a function of phase initiation

The previous analyses demonstrated that participants started tactile sampling in either phase of the cardiac cycle, but they preferentially finished sampling in diastole. This is consistent with the prediction that we are more sensitive to tactile information in diastole than systole. To explore this further we tested whether tactile sampling would be dependent upon the phase of the cardiac cycle in which the touch was initiated. Specifically, we tested the prediction that if the tactile sampling was initiated in systole the duration of the touch would be greater to counter the reduced perceptual tactile sensitivity in systole. To address this, we calculated the length of time that participants spent holding their touch when the touches were initiated in systole or diastole. This duration was calculated for each level of difficulty, generating a 2×7 factorial design with factors cardiac phase and task difficulty.

The rmANOVA demonstrated a significant main effect of task difficulty (*F*(1.802, 81.073) = 26.292, *p* < 0.0001, *η*_*p*_^*2*^ = 0.369), as well as a main effect of cardiac phase (*F*(1, 45) = 17.382, *p* = 0.0001, *η*_*p*_^*2*^ = 0.279). As predicted, the length of holding time was greater when participants started to touch in systole 1143ms (SD = 486.6) compared to diastole 1093 (SD = 443.4; Figure 4A). The interaction between cardiac phase and difficulty, after sphericity corrections, was not significant *F*(3.917, 176.267) = 2.426, *p* = 0.051, *η*_*p*_^*2*^ = 0.051. The equivalent analysis for the control flat stimulus revealed no significant differences in touch duration when the touch was initiated in systole (M = 708, SD = 441.7) and diastole (M = 717, SD = 429.4), *Z* = 461.5, *p* = 0.531, *r* = -0.108, 95% CI mean difference [-25.25, 13.50]; Figure 4B. Overall, these results demonstrate that the duration of participants’ touch varied as a function of the phase of the cardiac cycle in which they initiated the touch. Furthermore, and importantly, this modulation was only observed when tactile discrimination was required.

**Figure 4.**
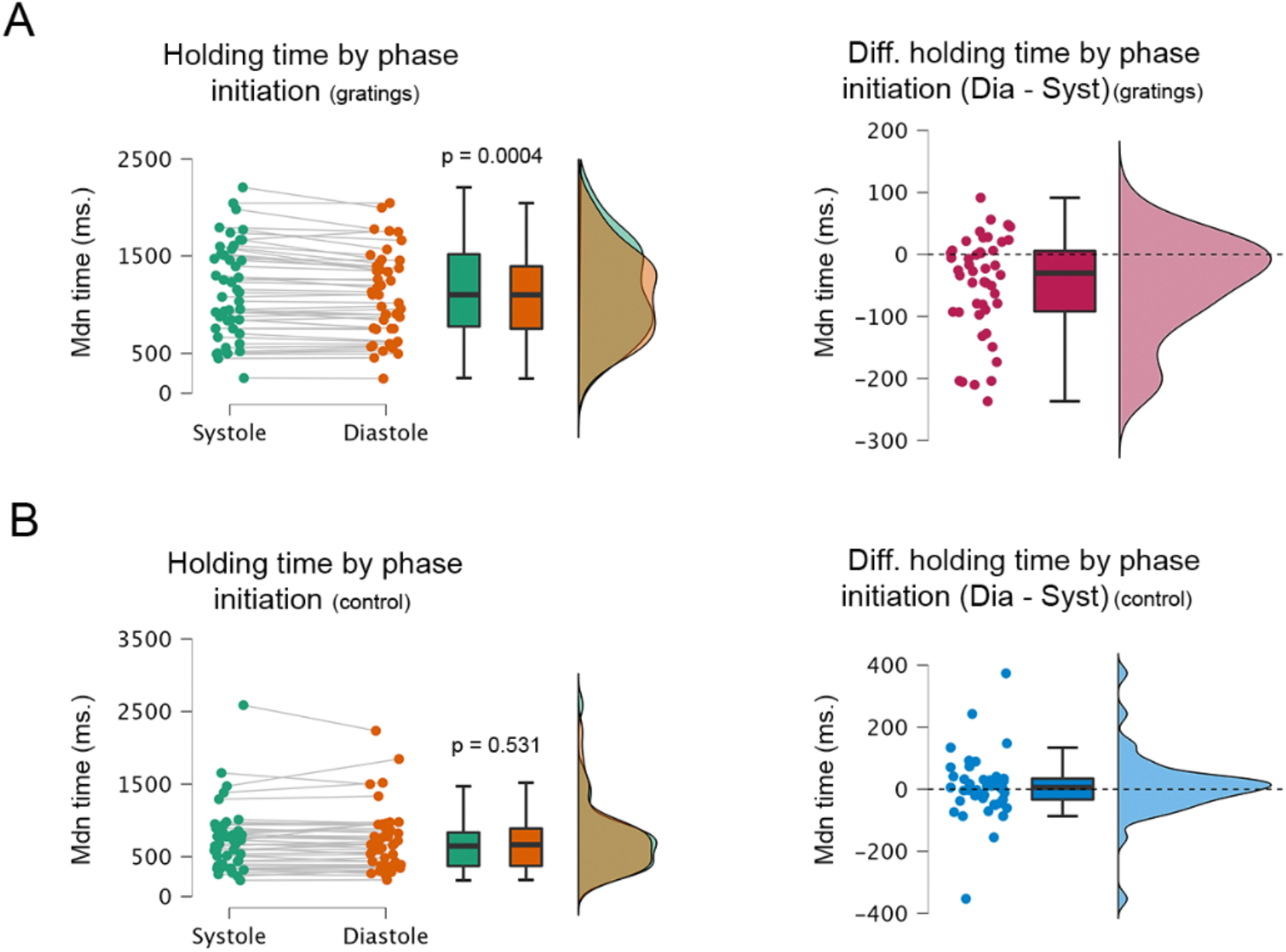
Sensing duration as a function of the initial cardiac phase. **(A)** The length of holding time was greater when participants started to touch in systole vs. diastole (*p* = 0.0004; M_diastole_ = 1143, M_systole_ = 1093). **(B)** Conversely, for the control flat stimulus the length of holding time was similar when participants started to touch in systole and diastole (*p* = 0.531; M_diastole_ = 708, M_systole_ = 717). Right panels show the difference between the duration of stationary holds initiated in diastole and systole (i.e., for each participant, the holding time of touches started in diastole minus touches started in systole).

#### Time sensing by phase initiation and touch variability

In the previous analysis, we demonstrated that the duration of the touch was dependent on the phase of the cardiac cycle in which the touch was initiated, touching for longer when the touch was initiated in systole. In this task, participants were free to initiate touching and to touch the grating for as long as they wished. Of interest is whether participants differed in their touching behaviour. Specifically, whether those who touched for longer did so to overcome the reduced tactile sensitivity in systole. In an initial qualitative analysis, we investigated whether there were any systematic differences in participants’ touching behaviour, specifically the duration that participants touched the gratings. We found that there was considerable variability in how long participants touched the gratings during the task (Figure 4B). Some participants held for relatively short periods of time and did not modulate their touching duration with task difficulty. In contrast, some participants held for longer periods and increased their touching time with task difficulty. To capture this between-subject variance, we calculated for each participant the standard deviation of how long they held the sensor across all difficulty levels. Note that this value is independent of difficulty level. Next, we ran again the previous analysis but with this variable as a between-subject covariate.

When adjusting the results by this covariate, the main effect of task difficulty in the rmANOVA was still significant (*F*(2.909, 127.993) = 3.070, *p* = 0.032, *η*_*p*_^*2*^ = 0.065), and the effect of cardiac phase and its interaction with task difficulty were not significant (*F*(1, 44) = 0.923, *p* = 0.342, *η*_*p*_^*2*^ = 0.021) and (*F*(4.089, 179.904) = 0.0896, *p* = 0.469, *η*_*p*_^*2*^ = 0.020). In addition, there was a significant interaction between difficulty and touch variability (*F*(2.909, 127.993) = 41.706, *p* < 0.0001, *η*_*p*_^*2*^ = 0.487), a significant interaction between cardiac phase and touch variability (*F*(1, 44) = 17.728, *p* = 0.0001, *η*_*p*_^*2*^ = 0.287), and a significant interaction between task difficulty, cardiac phase and touch variability (*F*(4.089, 179.904) = 4.239, *p* = 0.002, *η*_*p*_^*2*^ = 0.088). In addition, the correlation between participants’ proportion of correct responses and the variability of their holding time was also significant (Spearman’s rho = 0.401, *p* = 0.006; Supp. Figure 1) demonstrating that participants who varied the duration of the stationary holds in the experiment were also those with greater tactile discrimination performance,

Overall, these results demonstrate that the duration of the participants’ touch varied as a function of the cardiac phase in which they initiated it. Touches initiated in systole were held for longer periods of time, consistent with the hypothesis that tactile sensitivity is reduced in systole and participants had to hold for longer when touching in systole to overcome this. This effect was driven by those participants who varied the length of holding time in the experiment. This effect was most marked when these participants were touching the most difficult gratings to discriminate (Figure 5A,C,D).

**Figure 5.**
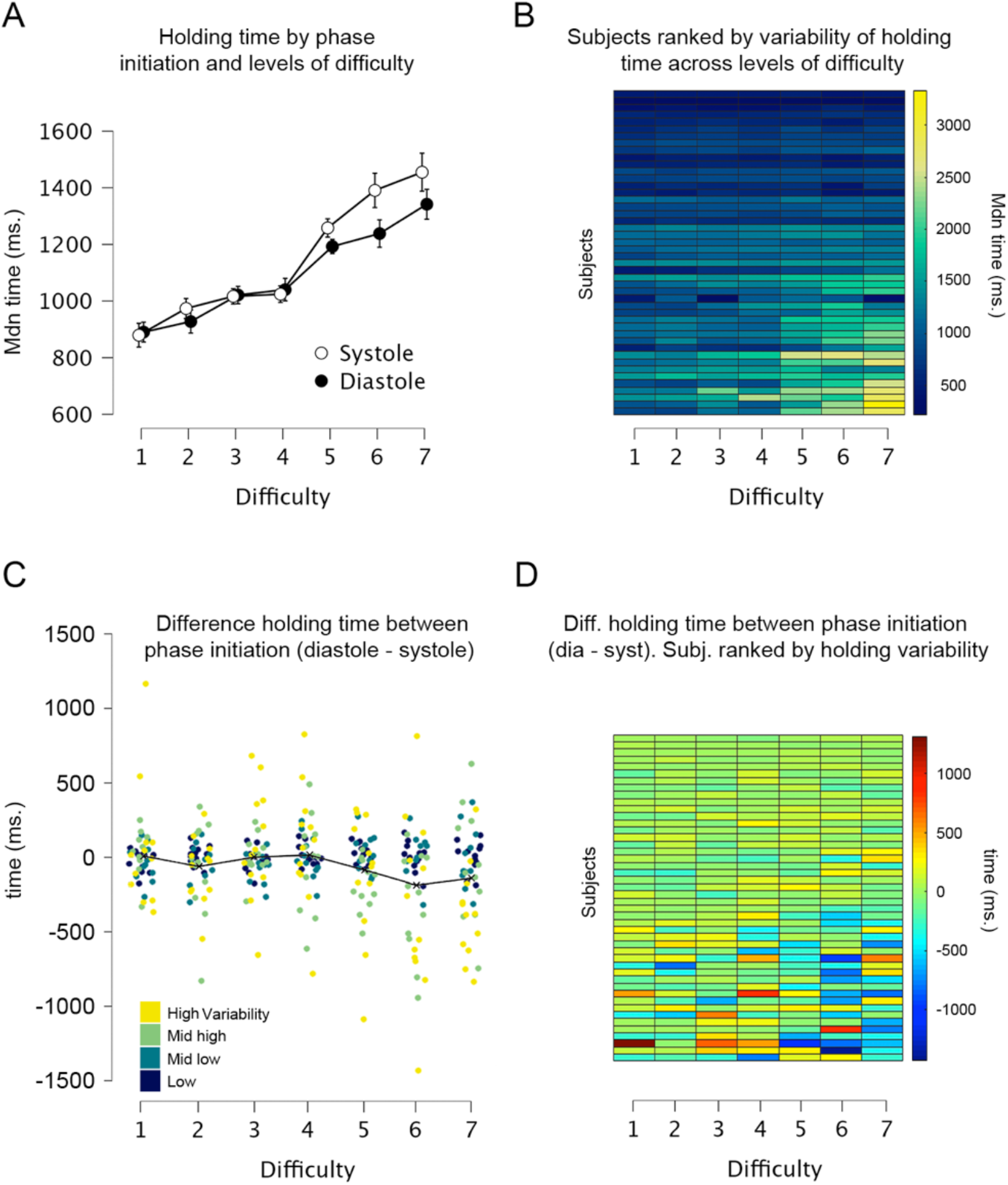
Sensing duration as a function of initial cardiac phase and touch variability. **(A)** Across all levels of difficulty (1 easiest, 7 most difficult), duration of stationary holds when participants started touching the gratings in each cardiac phase. **(B)** Participants’ duration of stationary holds varied across the different levels of difficulty (variability of holding time in the task). The heatmap depicts all participants ranked as a function of touch variability. Each row depicts one participant and each column one level of difficulty. Participants at the top varied the least the duration of their stationary holds. **(C)** Difference between the duration of stationary holds initiated in diastole and systole (i.e., for each participant and level of difficulty, holding time of touches started in diastole minus started in systole); dots’ colouration denotes participants’ variability. Participants who varied the duration of their stationary holds, spent less time holding their touch when they started touching in diastole across the most difficult levels (task difficulty x cardiac phase *touch variability: *p* = 0.002). **(D)** Data from panel C depicted as a heatmap with participants ranked as a function of touch variability.

#### Cardiac interbeat interval as a function of starting touch

The previous analysis demonstrated that participants modulated the duration of the touch in a manner that was consistent with the reduced tactile sensitivity during systole. Given that tactile sensitivity is known to be reduced during systole another mechanism that could facilitate tactile perception would be for the participants to increase the duration of the cardiac cycle in diastole. To test this for each touch, we calculated the interbeat interval (i.e., IBI) for the cardiac cycle in which participants initiated the touch, as well as the two preceding and subsequent interbeat intervals. There was a significant main effect of heartbeat position on the duration of the IBIs (*F*(2.062, 92.788) = 11.65, *p* < 0.0001, *η*_*p*_^*2*^ = 0.206.) Post hoc comparisons showed that the duration of the IBI in the starting touch (M = 797.8, SD = 111.7) was significantly longer than in the remaining heartbeats (all *p* < 0.006, all *d >* 0.502; M_-2_ = 784.2, M_-1_ = 789.9, M_+1_ = 789.8, M_+2_ = 783.7; adjusted for multiple comparisons).

For the control flat stimulus, the results also show a significant main effect of heartbeat position on the duration of the IBIs *F*(2.595, 114.201) = 10.75, *p* < 0.0001, *η*_*p*_^*2*^ = 0.196. Post hoc comparisons revealed that the duration of the IBI in the starting touch (M = 821.4, SD = 118.6) was significantly longer than in the two previous heartbeats (all *p* < 0.015, all *d >* 0.473; M_-2_ = 802.5, M_-1_ = 810.6). Also, the IBI of the most preceding heartbeat was shorter than the two latter IBIs (all *p* < 0.016, all *d > -*0.443, adjusted for multiple comparisons). See Supp. Table 3-6 for all Post hoc comparisons.

Lastly, we examined whether these differences were driven by changes in the length of the diastole or systole phases of the cardiac cycle. For both, the grating and control flat stimuli, the systolic duration of preceding and successive heartbeats relative to the staring touch did not vary significantly (*p* > 0.235, *η*_*p*_^*2*^ < 0.032). For the diastole phase, the analyses showed a significant main effect of heartbeat position (*p* < 0.0001, *η*_*p*_^*2*^ > 0.200) and the Post hoc analysis showed similar differences to those observed between the interbeat intervals of the positions relative to the staring touch (Fig. 6A,B, Supp. Tables 3-6). These results confirm the lengthening of time between consecutive heartbeats (cardiac deceleration) in anticipation of stimulus or in reaction to a salient stimulus (e.g., Lacey and Lacey, 1977; Sandman et al., 1977), which is typically followed by acceleration after the behavioural response (e.g., Coles and Duncan-Johnson, 1975; Park et al., 2014). This is also consistent with the hypothesis that participants increased the proportion of time in diastole to increase tactile perceptual sensitivity. This is also confirmed in a supplementary analysis where we observed that the duration of subjects’ touch as a function of the cardiac phase in which they initiated it was significant when adding the covariate heartbeat deacceleration; see Supp. Table 7.

**Figure 6.**
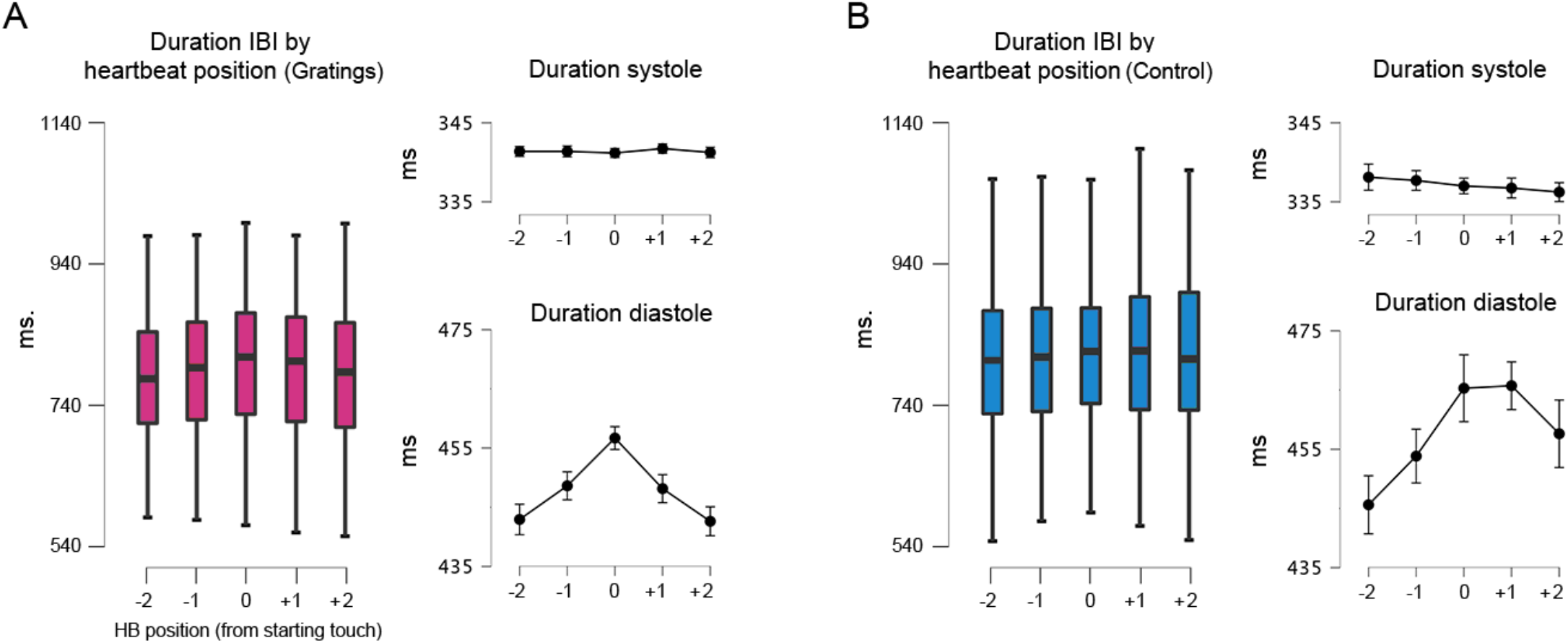
Interbeat interval before, during, and after the touch of the stimuli. **(A)** For the grating stimuli, the interbeat interval (IBI) of the heartbeat where the touch was initiated (0 in the x-axis) was significantly longer than the preceding and succeeding heartbeats (*p* < 0.0001). Right panels: with similar statistical effects, this difference was driven by the elongation of the diastole phase of the cardiac cycle. **(B)** For the control flat stimulus, the interbeat interval (IBI) of the heartbeat where the touch was initiated was significantly longer than in the two preceding heartbeats (*p* < 0.015). Also, the IBI of the earliest heartbeat differed from the two later ones and the IBI preceding the heartbeat differed from the later one (relative to 0). Right panels: with similar statistical effects, this difference was also driven by the elongation of the diastole phase of the cardiac cycle; all p-values adjusted using the Holm-Bonferroni method.

## Discussion

The effect of the cardiac phases upon stimulus processing has previously been observed by time-locking the presentation of stimuli to the cardiac phase (e.g., Al et al., 2021; Critchley & Garfinkel, 2018; Garfinkel et al., 2020). However, in our everyday experiences, sensory information is not encountered in such a phase-locked manner. Here we tested the following hypothesis: If the phase of the cardiac cycle is an important modulator of perception and cognition, as previously proposed, it should modulate the way in which we freely and actively seek information in the world. To test this, we recorded the ECGs from human participants while they performed a tactile discrimination task of grating orientation. Importantly, they freely initiated, held, and ended the sensing of the stimuli.

From an interoceptive inference framework, studies in passive sensing have shown that tactile sensitivity is reduced during the systole phase of the cardiac cycle (Al, Iliopoulos, et al., 2021; Grund et al., 2022; Motyka et al., 2019). Hence, to avoid this drop in sensitivity, participants could circumvent the sampling of information in systole. This is precisely what the results of the present work suggest. The duration of subjects’ touch (stationary holds) varied as a function of the cardiac phase in which they initiated it. Touches initiated in the systole phase of the cardiac cycle (systole) were held for longer periods of time. This is consistent with the idea that participants actively sought to sample longer when they initiated their touch in a period with reduced perceptual tactile sensitivity. Furthermore, we demonstrated, that this effect was driven by those participants who varied the length of their touches as a function of task difficulty. In addition, this modulation in the touching time was associated with improved tactile discrimination performance. Conversely, while touches in the control condition were coupled to the cardiac cycle, their length was not modulated as a function of when in the cycle these were initiated. This latter condition only required movement, which seems facilitated during systole (Al et al., 2021; Galvez-Pol et al., 2020; Konttinen et al., 2003; Kunzendorf et al., 2019; Ohl et al., 2016; Palser et al. 2021). In addition, we showed that after touch initation, participants had a significant deceleration of their heart rate driven by an increased time in diastole. This is also consistent with the idea that participants increased their active sampling in the diastole phase by increasing the time spent in this period.

### Coupling between active sensing and transient bodily cycles

Increasing evidence suggests that ever-fluctuating interoceptive signals exert a putative effect over stimulus processing (Azevedo et al., 2017; Azzalini et al., 2019, 2021; Garfinkel & Critchley, 2016; Rae et al., 2018). This work is often conceptualized within the predictive coding framework (Allen et al., 2019; Friston et al., 2017) to explain how the brain integrates interoceptive and exteroceptive information to optimise beliefs about the world. Considering this, prevailing models of perception and learning suggest that humans and animals do not only update their beliefs with incoming sensory evidence, but also seek information in a way that maximizes the reduction of uncertainty (Yang et al., 2016). This is reflected in the way humans and animals purposively seek information through the movement and control of the sensor apparatus (e.g., whiskers, eyes, fingers, antennae) in whatever manner best suits the task (Grant et al., 2014; Prescott et al., 2011; Yang et al., 2016).

The finding that participants’ touch in the control condition was coupled to the cardiac cycle, but that its length did not vary as a function of when in the cycle was initiated, is congruent to recent evidence showing that during simple hand movements, high corticospinal excitability and desynchronization of sensorimotor oscillations are observed in systole (Al et al., 2021). However, we reason that the fine use of the sensorium might entail alignment of behaviour and quiescent bodily cycles. Specifically, during the systole phase of the cardiac cycle, there are several physiological fluctuations. For instance, pressure sensors located in the carotid sinus, coronary arteries, and aortic arch (baroreceptors), detect changes in blood pressure and convey the strength and timing of heartbeats to the brainstem through the vagus nerve (Critchley & Harrison, 2013; Davos et al., 2002). Succeeding this, the closed-loop arterial baroreflex system, through at least three components (heart rate, vascular tone, and stroke volume), buffers blood pressure fluctuations (Vaschillo et al., 2011, 2012), and concomitant changes such as ballistic fluctuations of the ejected blood and arterial pulsations exert a direct effect in muscle activity (Birznieks et al., 2012; Fallon, 2004), as well as elicit small head and eye movements (Debener et al., 2010; Lauridsen et al., 2019). Altogether, these variations could interfere, compete, or block the allocation of attentional and/or representational resources for incoming sensory information. Thus, one possibility is that people actively seek more sensory information during the reduced presence of these interfering signals.

Early evidence for this account, comes from studies showing that elite rifle shooters pull the trigger during diastole, with beginners firing during either phase but with better results during diastole (Helin et al., 1987). It has been suggested that this effect is related to the mechanical movement caused by the heart’s contraction (Konttinen et al., 2003), which is also the principle suggested in the sampling of active visual information during silent cardiac activity (Galvez-Pol et al., 2020). Relatedly, an alignment between human participants and behaviour has been also shown in another facet of the cardiovascular system, i.e., respiration. Perl et al., (2019) showed the alignment of self-initiated cognitive tests with the beginning of the inspiratory phase, and recent work by Kluger et al., (2021) has shown respiration-locked performance in a near-threshold spatial detection task.

### Constraints of generality and future work

Here we consider the constraints of generality proposed by Simons et al., (2017). First, the type of tactile discrimination used here (gratings orientation) represents one of the vast spectra of haptic behaviours that people use in daily life (e.g., detection, discrimination and/or identification of stimulus’ position, texture, size, shape, etc). Second, while we isolated the sense of touch from other exteroceptive modalities such as vision, these and other modalities frequently work together at the sensor and neuronal levels (Banati et al., 2000; Galvez-Pol et al., 2020; Taylor-Clarke et al., 2002; Zhou & Fuster, 2000). Third, our stimuli were judged by young adults (18-35yo) recruited through a university subject pool. Thus, we expect the results to generalize to situations in which participants have to engage with similar stimuli similarly. Moreover, we believe the results reported here will be reproducible with young adults from similar subject pools serving as participants. However, we do not have enough evidence to state that our findings will occur outside of these settings, nor to state which changes could be observed when either the stimuli or type of task differ from those reported here. We have no reason to believe that the results depend on other characteristics of the participants and materials.

Future research could examine whether the functional alignment of the sensorium with quiescent periods of the cardiovascular system interacts with different levels of conscious interoceptive inferences. To this aim, further work could examine how active sensing unfolds in individuals that sense, interpret, and integrate to a different extent signals originating from within the body (e.g., interoceptive sensibility; e.g., Galvez-Pol et al., 2021; Garfinkel et al., 2015; Palser et al., 2018; Suksasilp & Garfinkel, 2022). Likewise, future work in active sensing may include the coregistration of neuronal and physiological recordings. This might include, for instance, heartbeat, motor-cortical, and/or somatosensory-evoked activity (e.g., Al et al., 2020; Galvez-Pol et al., 2018, 2020; Montoya et al., 1993; Park & Blanke, 2019). As described in Kluger et al., (2021), bodily-entrained fluctuations in neural activity may represent a mechanism for uniting neural and behavioural findings. While evidence for this account has been accumulating in the animal literature, specially between the respiratory cycle and behaviours (Kurnikova et al., 2017; Moore et al., 2013), the field of active sensing in humans with physiological measures remains largely uncharted. Here the inclusion of the cardiac cycle, an intrinsic oscillator with a faster cycle, might aid to understand the hierarchical organisation of neural oscillations (Lakatos et al., 2005) altogether with bodily rhythms such as those originating from the whole cardiovascular system.

## Acknowledgements

We would like to thank all those who participated in and helped to advance this study.

## Data and materials availability

The code and anonymised data are available in the following Open Science Framework repository: https://osf.io/d7x3g/

## Author contributions according to CRediT statement

A.G-P: Conceptualization, Investigation, Methodology, Software, Data Acquisition, Visualization, Formal Analysis, Writing original draft. P.V: Investigation, Visualization, Data Acquisition. J.V: Visualization, Formal Analysis, Writing - review & editing. J-M.K: Conceptualization, Investigation, Methodology, Software, Visualization, Formal Analysis, Writing - review & editing, Project administration, Funding acquisition.

