## Supplementary materials for "Active tactile discrimination is coupled with and modulated by the cardiac cycle"

**Supp. material 1. Instructions for participants.** In this experiment we will present to you small gratings with different widths (from narrow to wider gratings). These gratings have two possible orientations: i) horizontal or ii) vertical. Here we will randomly present to you one grating at the time (one per trial). Your task is to touch these gratings with your index finger and tell us whether the grating is oriented vertically or horizontally. In each trial, the experimenter will randomly select one of the gratings to-be-touched. Once the experimenter has placed the grating in position, she/he will place both hands on top of the device (this means that the grating is ready and secured). Then, feel free to touch the grating whenever you feel like, and for as long as you feel like. Yet do not overthink. For this, use your index finger and please produce a single tap ('touch-in'). Make sure you make firm contact with the grating. Once you touch, do not swing, swipe, or turn your finger (you should 'touch-in and touch-out' by just moving your finger up and down). Once you know the orientation, 'Touch-out' and only then tell us verbally the orientation of the grating (we will key your response). Please tell us the orientation even if you are uncertain. To get familiar with the task, you will do a practise block. Next, you will do 12 blocks of ~4 mins each. Do you have any questions?

**Supp. Table 1. Post Hoc comparisons - Proportion correct responses by gratings difficulty (1-7)**

| | | Mean Difference | SE | t | Cohen's d | $p_{holm}$ |
|---|---|---|---|---|---|---|
| 1 | 2 | 0.009 | 0.020 | 0.462 | 0.068 | 1.000 |
| | 3 | 0.025 | 0.020 | 1.238 | 0.183 | 1.000 |
| | 4 | 0.051 | 0.020 | 2.522 | 0.372 | 0.086 |
| | 5 | 0.165 | 0.020 | 8.169 | 1.204 | 1.330e−13*** |
| | 6 | 0.386 | 0.020 | 19.142 | 2.822 | 6.497e−51*** |
| | 7 | 0.405 | 0.020 | 20.045 | 2.955 | 4.742e−54*** |
| 2 | 3 | 0.016 | 0.020 | 0.777 | 0.115 | 1.000 |
| | 4 | 0.042 | 0.020 | 2.061 | 0.304 | 0.242 |
| | 5 | 0.156 | 0.020 | 7.707 | 1.136 | 2.469e−12*** |
| | 6 | 0.377 | 0.020 | 18.680 | 2.754 | 2.512e−49*** |
| | 7 | 0.395 | 0.020 | 19.583 | 2.887 | 1.895e−52*** |
| 3 | 4 | 0.026 | 0.020 | 1.284 | 0.189 | 1.000 |
| | 5 | 0.140 | 0.020 | 6.930 | 1.022 | 2.774e−10*** |
| | 6 | 0.361 | 0.020 | 17.903 | 2.640 | 1.372e−46*** |
| | 7 | 0.380 | 0.020 | 18.806 | 2.773 | 9.495e−50*** |
| 4 | 5 | 0.114 | 0.020 | 5.646 | 0.833 | 3.317e−7*** |
| | 6 | 0.336 | 0.020 | 16.620 | 2.450 | 4.669e−42*** |
| | 7 | 0.354 | 0.020 | 17.522 | 2.584 | 2.948e−45*** |
| 5 | 6 | 0.222 | 0.020 | 10.973 | 1.618 | 2.486e−22*** |
| | 7 | 0.240 | 0.020 | 11.876 | 1.751 | 2.447e−25*** |
| 6 | 7 | 0.018 | 0.020 | 0.903 | 0.133 | 1.000 |

*** $p < .001$

*Note.* Cohen's d does not correct for multiple comparisons.

*Note.* P−value adjusted for comparing a family of 21

**Supp. Table 2. Post Hoc comparisons - Proportion Mdn holding times by gratings difficulty (1-7)**

| | | Mean Difference | SE | t | Cohen's d | $p_{holm}$ |
|---|---|---|---|---|---|---|
| 1 | 2 | −80.87 | 55.25 | −1.464 | −0.216 | 0.722 |
| | 3 | −128.08 | 55.25 | −2.318 | −0.342 | 0.148 |
| | 4 | −147.25 | 55.25 | −2.665 | −0.393 | 0.065 |
| | 5 | −354.57 | 55.25 | −6.418 | −0.946 | 9.891e−9*** |
| | 6 | −414.30 | 55.25 | −7.499 | −1.106 | 1.763e−11*** |
| | 7 | −527.49 | 55.25 | −9.548 | −1.408 | 1.766e−17*** |
| 2 | 3 | −47.21 | 55.25 | −0.854 | −0.126 | 0.922 |
| | 4 | −66.38 | 55.25 | −1.202 | −0.177 | 0.922 |
| | 5 | −273.70 | 55.25 | −4.954 | −0.730 | 1.669e−5*** |
| | 6 | −333.43 | 55.25 | −6.035 | −0.890 | 7.827e−8*** |
| | 7 | −446.62 | 55.25 | −8.084 | −1.192 | 4.239e−13*** |
| 3 | 4 | −19.17 | 55.25 | −0.347 | −0.051 | 0.922 |
| | 5 | −226.49 | 55.25 | −4.100 | −0.604 | 6.029e−4*** |
| | 6 | −286.23 | 55.25 | −5.181 | −0.764 | 6.059e−6*** |
| | 7 | −399.41 | 55.25 | −7.230 | −1.066 | 8.981e−11*** |
| 4 | 5 | −207.32 | 55.25 | −3.752 | −0.553 | 0.002** |
| | 6 | −267.05 | 55.25 | −4.834 | −0.713 | 2.701e−5*** |
| | 7 | −380.24 | 55.25 | −6.882 | −1.015 | 6.976e−10*** |
| 5 | 6 | −59.74 | 55.25 | −1.081 | −0.159 | 0.922 |
| | 7 | −172.92 | 55.25 | −3.130 | −0.461 | 0.017* |
| 6 | 7 | −113.18 | 55.25 | −2.049 | −0.302 | 0.249 |

*Note.* P-value adjusted for comparing a family of 21
*Note.* Cohen's d does not correct for multiple comparisons.
* $p < .05$, ** $p < .01$, *** $p < .001$

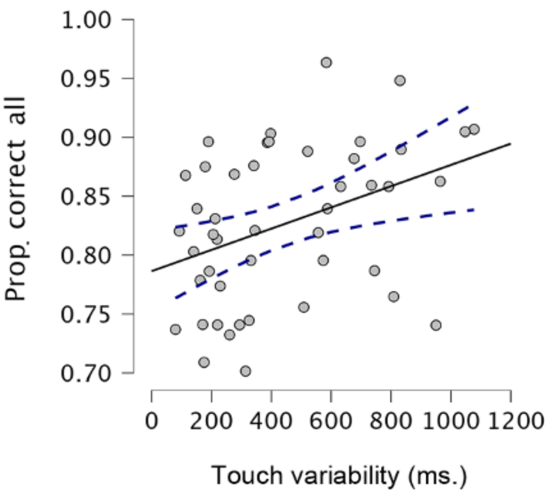


**Supp Fig. 1. Proportion of correct responses with touch variability.** The overall proportion of participants' correct responses correlated positively (Spearman's rho = 0.401, $p$ = 0.006) with participants' touch variability (i.e., as the standard deviation of how long in milliseconds they held the sensor across all levels of difficulty).

**Supp. Table 3. Post Hoc tests – Duration IBIs by heartbeat position (relative to touch entailing heartbeat) – Gratings**

| | | Mean Difference | SE | t | Cohen's d | $p_{holm}$ |
|---|---|---|---|---|---|---|
| −2 | −1 | −5.754 | 2.363 | −2.434 | −0.359 | 0.064 |
| | 0 | −13.641 | 2.363 | −5.772 | −0.851 | 3.031e−7*** |
| | +1 | −5.584 | 2.363 | −2.363 | −0.348 | 0.064 |
| | +2 | 0.443 | 2.363 | 0.188 | 0.028 | 1.000 |
| −1 | 0 | −7.888 | 2.363 | −3.338 | −0.492 | 0.007** |
| | +1 | 0.169 | 2.363 | 0.072 | 0.011 | 1.000 |
| | +2 | 6.197 | 2.363 | 2.622 | 0.387 | 0.057 |
| 0 | +1 | 8.057 | 2.363 | 3.409 | 0.503 | 0.006** |
| | +2 | 14.085 | 2.363 | 5.960 | 0.879 | 1.303e−7*** |
| +1 | +2 | 6.028 | 2.363 | 2.551 | 0.376 | 0.058 |

*Note.* P-value adjusted for comparing a family of 10
*Note.* Cohen's d does not correct for multiple comparisons.
* p < .05, ** p < .01, *** p < .001

**Supp. Table 4. Post Hoc tests – Duration IBIs by heartbeat position - Flat control stimulus**

| | | Mean Difference | SE | t | Cohen's d | $p_{holm}$ |
|---|---|---|---|---|---|---|
| −2 | −1 | −8.091 | 3.407 | −2.375 | −0.354 | 0.068 |
| | 0 | −18.917 | 3.407 | −5.553 | −0.828 | 1.022e−6*** |
| | +1 | −18.564 | 3.407 | −5.449 | −0.812 | 1.519e−6*** |
| | +2 | −10.350 | 3.407 | −3.038 | −0.453 | 0.017* |
| −1 | 0 | −10.827 | 3.407 | −3.178 | −0.474 | 0.014* |
| | +1 | −10.474 | 3.407 | −3.074 | −0.458 | 0.017* |
| | +2 | −2.259 | 3.407 | −0.663 | −0.099 | 1.000 |
| 0 | +1 | 0.353 | 3.407 | 0.104 | 0.015 | 1.000 |
| | +2 | 8.567 | 3.407 | 2.515 | 0.375 | 0.064 |
| +1 | +2 | 8.214 | 3.407 | 2.411 | 0.359 | 0.068 |

*Note.* P-value adjusted for comparing a family of 10

**Supp. Table 5. Post Hoc tests – Duration diastole by heartbeat position - Gratings**

| | | Mean Difference | SE | t | Cohen's d | $p_{holm}$ |
|---|---|---|---|---|---|---|
| −2 | −1 | −8.194 | 3.465 | −2.365 | −0.353 | 0.096 |
| | 0 | −19.420 | 3.465 | −5.604 | −0.835 | 7.165e−7*** |
| | +1 | −19.853 | 3.465 | −5.729 | −0.854 | 4.297e−7*** |
| | +2 | −11.936 | 3.465 | −3.444 | −0.513 | 0.006** |
| −1 | 0 | −11.225 | 3.465 | −3.239 | −0.483 | 0.009** |
| | +1 | −11.658 | 3.465 | −3.364 | −0.502 | 0.007** |
| | +2 | −3.741 | 3.465 | −1.080 | −0.161 | 0.564 |
| 0 | +1 | −0.433 | 3.465 | −0.125 | −0.019 | 0.901 |
| | +2 | 7.484 | 3.465 | 2.160 | 0.322 | 0.096 |
| +1 | +2 | 7.917 | 3.465 | 2.285 | 0.341 | 0.096 |

*Note.* P-value adjusted for comparing a family of 10

**Supp. Table 6. Post Hoc tests – Duration diastole by heartbeat position - Flat control stimulus**

| | | Mean Difference | SE | t | Cohen's d | $p_{holm}$ |
|---|---|---|---|---|---|---|
| −2 | −1 | −5.747 | 2.418 | −2.377 | −0.350 | 0.093 |
| | 0 | −13.860 | 2.418 | −5.732 | −0.845 | 3.699e−7*** |
| | +1 | −5.218 | 2.418 | −2.158 | −0.318 | 0.097 |
| | +2 | 0.309 | 2.418 | 0.128 | 0.019 | 1.000 |
| −1 | 0 | −8.113 | 2.418 | −3.355 | −0.495 | 0.007** |
| | +1 | 0.529 | 2.418 | 0.219 | 0.032 | 1.000 |
| | +2 | 6.056 | 2.418 | 2.505 | 0.369 | 0.079 |
| 0 | +1 | 8.642 | 2.418 | 3.574 | 0.527 | 0.004** |
| | +2 | 14.169 | 2.418 | 5.860 | 0.864 | 2.163e−7*** |
| +1 | +2 | 5.527 | 2.418 | 2.286 | 0.337 | 0.094 |

*Note.* P-value adjusted for comparing a family of 10

**Supp. Table 7. Mdn holding times by cardiac phase x level of difficulty * heartbeat deacceleration**

Within Subjects Effects

| Cases | Sum of Squares | df | Mean Square | F | p | $\eta^2_p$ |
|---|---|---|---|---|---|---|
| Level difficulty | 8.735e+6[a] | 6[a] | 1.456e+6[a] | 10.806[a] | 9.484e−11[a] | 0.197 |
| Level difficulty * HB0-2_All_lvls | 384310.07[a] | 6[a] | 64051.68[a] | 0.475[a] | 0.826[a] | 0.011 |
| Residuals | 3.557e+7 | 264 | 134729.40 | | | |
| Cardiac Phase | 210536.62 | 1 | 210536.62 | 9.116 | 0.004 | 0.172 |
| Cardiac Phase * HB0-2_All_lvls | 62.87 | 1 | 62.87 | 0.003 | 0.959 | 6.187e−5 |
| Residuals | 1.016e+6 | 44 | 23095.32 | | | |
| Level difficulty * Cardiac Phase | 514930.56[a] | 6[a] | 85821.76[a] | 2.276[a] | 0.037[a] | 0.049 |
| Level difficulty * Cardiac Phase * HB0-2_All_lvls | 1.039e+6[a] | 6[a] | 173236.96[a] | 4.594[a] | 1.864e−4[a] | 0.095 |
| Residuals | 9.955e+6 | 264 | 37709.54 | | | |

*Note*. Duration of subjects' touch as a function of the cardiac phase in which they initiated it (Cardiac phase) interacted significantly with task difficulty (Level of difficulty) when the covariate heartbeat deacceleration was added (i.e., HB0-2_All_lvls). Deacceleration was computed by subtracting the duration of the heartbeat entailing the starting touch minus that of the second heartbeat before the starting touch (i.e., IBI entailing the touch in ms minus IBI of the second heartbeat before touch). Level of difficulty x Cardiac Phase * HBO-2_All_lvls, $p = 0.0004$.
